# Hypoxia induces cytotoxicity and suppresses cytokine production by CD8□ T cells in cutaneous leishmaniasis

**DOI:** 10.64898/2026.07.30.741849

**Authors:** Erin A. Fowler, Olivia L. Schneider, Faaiza Saif, Fernanda O. Novais

## Abstract

Cutaneous leishmaniasis is characterized by chronic inflammatory skin lesions in which CD8□ T cells exhibit paradoxical functions. While IFN-γ-producing CD8□ T cells contribute to the development of protective immunity in the draining lymph node, CD8□ T cells recruited to the infected skin lose their ability to produce IFN-γ and instead acquire cytotoxic functions that exacerbate tissue damage. We previously demonstrated that the hypoxic microenvironment of leishmanial lesions promotes CD8□ T cell cytotoxicity through induction of Blimp-1. Whether hypoxia also suppresses protective CD8□ T cell functions, however, is unknown. Here, we show that hypoxia simultaneously suppresses production of the protective cytokines IFN-γ and TNF-α while enhancing expression of granzyme B and perforin in activated CD8□ T cells. In vitro, HIF-1α, but not HIF-2α, was required for hypoxia-induced expression of granzyme B, perforin, and Blimp-1, whereas suppression of IFN-γ and TNF-α occurred independently of HIF signaling, indicating that distinct oxygen-related pathways regulate pathogenic and protective CD8□ T cell functions. Hypoxia also increased expression of multiple inhibitory receptors on CD8□ T cells, although lesional CD8□ T cells lacked expression of the terminal exhaustion-associated transcription factor TOX, suggesting that hypoxia promotes an inhibitory phenotype distinct from terminal exhaustion. Finally, adoptive transfer studies demonstrated that in vivo both HIF-1α and HIF-2α expression in CD8□ T cells contributed to immunopathology during cutaneous leishmaniasis. Together, these findings identify hypoxia as a key regulator that functionally reprograms CD8□ T cells by promoting pathogenic cytotoxicity while suppressing protective cytokine production within lesions.

## Introduction

Cutaneous leishmaniasis is a chronic inflammatory disease caused by protozoan parasites of the genus *Leishmania* that is characterized by chronic skin ulcers. Disease severity is influenced by both parasite species and the host immune response. Protective immunity depends on IFN-γ and TNF-α-driven activation of infected macrophages, primarily mediated by CD4□ Th1 cells, which promote parasite killing and lesion resolution (1). CD8□ T cells also contribute to host defense during the early stages of infection by producing IFN-γ in the draining lymph node (dLN), where they support the development of protective Th1 responses. However, once recruited into the infected skin, CD8□ T cells undergo a phenotypic switch. Rather than producing IFN-γ, they acquire a cytotoxic phenotype characterized by granzyme B and perforin expression that exacerbates tissue damage without controlling the parasite (2–4). Understanding how the tissue microenvironment drives this transition from protective to pathogenic CD8□ T cell function remains unknown.

One critical characteristic of chronically inflamed tissues is reduced oxygen availability. Hypoxia develops within tumors and sites of inflammation as oxygen demand exceeds supply, and is a critical regulator of immune cell function. Beyond influencing cellular metabolism, hypoxia alters transcriptional and metabolic programs that influence cytokine production, cytotoxicity, survival, and tissue adaptation (5, 6). We and others have previously demonstrated that *Leishmania*-infected lesions in both mice and humans are hypoxic and that exposure of CD8□ T cells to hypoxia promotes expression of granzyme B and perforin, resulting in increased immunopathology (7–9). Mechanistically, we identified hypoxic induction of the transcription factor Blimp-1 as a critical regulator of granzyme B expression, whereas perforin induction happened independently of Blimp-1 (8, 9). These findings established hypoxia as a key driver of pathogenic CD8□ T cell differentiation but also raised the question of whether hypoxia simply enhances cytotoxicity or also suppresses CD8□ T cell protective responses.

Cellular adaptation to hypoxia is mediated primarily through stabilization of the hypoxia-inducible factor (HIF) family of transcription factors (6). HIF-1α and HIF-2α regulate overlapping but distinct transcriptional programs controlling metabolism, inflammation, survival, and cellular adaptation to reduced oxygen tension (10–17). In T cells, HIF signaling has been implicated in multiple aspects of effector differentiation, including cytotoxic function, metabolism, migration, and persistence (18–27). Despite these advances, the contribution of individual HIF isoforms to CD8□ T cell function during chronic parasitic infection is untested. In particular, while hypoxia has consistently been linked to enhanced cytotoxicity, its effects on the production of protective type 1 cytokines remain variable across disease models, with studies reporting increased, decreased, or unchanged IFN-γ production depending on the context (18–20, 24, 27–35). Whether hypoxia suppresses protective cytokine production during cutaneous leishmaniasis, and whether this process is regulated through HIF-dependent pathways, is unknown.

The loss of IFN-γ production by lesional CD8□ T cells also raises the possibility that chronic hypoxia promotes inhibitory programs that limit protective function. In tumors and chronic viral infections, prolonged hypoxia enhances expression of multiple inhibitory receptors and has been implicated in the development of dysfunctional T cell states (18–20, 23, 26, 33, 36–41). Similarly, CD8□ T cells isolated from cutaneous leishmaniasis lesions express elevated levels of inhibitory receptors, although whether these changes reflect terminal exhaustion remains unclear (8, 42–44).

Here, we investigated how hypoxia reshapes CD8□ T cell function during cutaneous leishmaniasis. We tested whether hypoxia simultaneously promotes pathogenic cytotoxicity while suppressing protective type 1 cytokine production, determined the need for HIF-1α and HIF-2α for these processes, and examined whether hypoxia induces checkpoint molecules associated with exhausted CD8□ T cell responses. Together, our findings identify hypoxia as a central regulator that functionally and spatially segregates protective and pathogenic CD8□ T cell effector functions during cutaneous leishmaniasis.

## Results

### Hypoxia promotes pathogenic effector function while suppressing protective cytokine production by CD8^+^ T cells

Our previous studies demonstrated that the hypoxic microenvironment of *Leishmania*-infected lesions promotes expression of granzyme B and perforin by CD8□ T cells, thereby enhancing tissue pathology (8, 9). However, whether hypoxia also contributes to the loss of protective CD8□ T cell function observed within lesions is unknown. Because lesional CD8□ T cells fail to produce IFN-γ despite retaining cytotoxic activity (3), we first asked whether hypoxia concurrently regulates these effector programs. To address this question, splenocytes or draining lymph node (dLN) cells from *Leishmania*-infected mice were stimulated with plate-bound anti-CD3 and soluble anti-CD28 under normoxic (21% O□) or hypoxic (1% O□) conditions for 48 hours. During the last 4 hours, cells also received PMA, ionomycin, and brefeldin A. Consistent with our previous reports (8, 9), hypoxia markedly increased expression of granzyme B and perforin by antigen-experienced (CD44^high^) CD8□ T cells (Fig. 1A-B). In contrast, hypoxia significantly reduced production of both IFN-γ and TNF-α (Fig. 1C-D), closely resembling the phenotype of lesional CD8□ T cells observed during infection (4). Together, these findings demonstrate that hypoxia not only enhances cytotoxic differentiation but functionally reprograms CD8□ T cells by promoting pathogenic effector functions while suppressing protective cytokine production.

**Figure 1.**
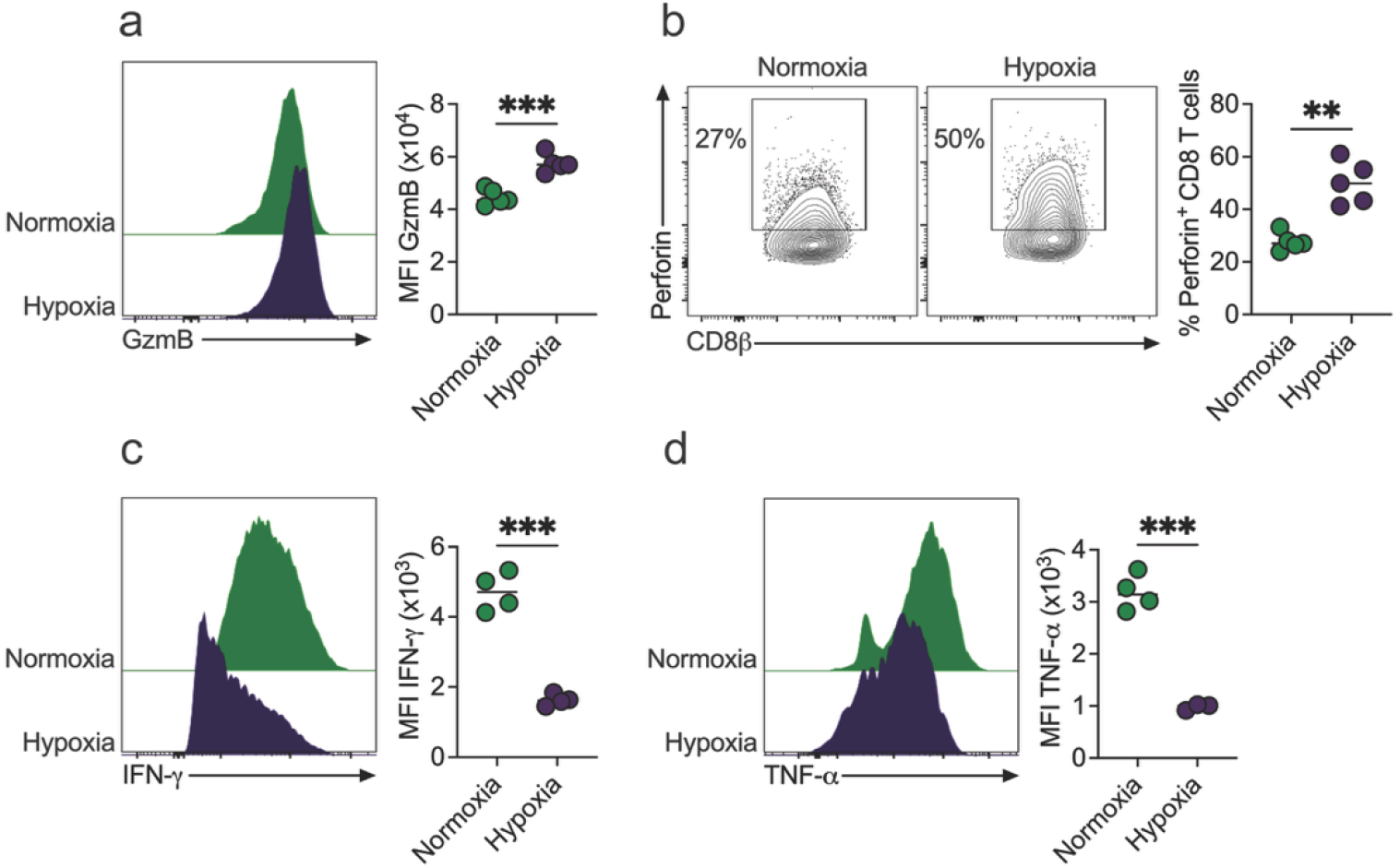
Hypoxia promotes cytotoxicity and suppresses type 1 cytokine production in CD8^+^ T cells. (a and. **b)** dLN single cell suspensions of *Leishmania major-*infected C57BL/6 mice were incubated in normoxia (21% O_2_) or hypoxia (1% O_2_) for 48 hours with anti-CD3 and anti-CD28 stimulation, and then CD8^+^ T cells were analyzed by flow cytometry for **(a)** mean fluorescence intensity (MFI) of granzyme B (GzmB) and **(b)** frequency of perforin expression. **(c and d)** dLN single cell suspensions of *L. major-*infected C57BL/6 mice were incubated in normoxia (21% O_2_) or hypoxia (1% O_2_) for 48 hours with anti-CD3 and anti-CD28 stimulation. For the last four hours of incubation, cells were stimulated with PMA (phorbol 12-myristate 13-acetate), ionomycin, and brefeldin A, and then CD8^+^ T cells were analyzed by flow cytometry for MFI of **(c)** IFN-γ and **(d)** TNF-α. **(a and b)** Representative contour plots, histograms, and scatter dot plots showing individual mice are representative of more than 4 experiments with at least 3 mice per experiment. **(c and d)** Representative contour plots, histograms, and scatter dot plots showing individual mice are representative of 2 experiments with at least 3 mice per experiment. Gating strategy: live, single cells, CD90, CD8β^+^, CD44^high^. ∗∗P ≤ .01 and ∗∗∗P ≤ .001 by 2-tailed Student’s t test.

### Distinct hypoxia-sensitive pathways regulate cytotoxic differentiation and type 1 cytokine production

The dual regulation of cytotoxic molecules and type 1 cytokines suggested that hypoxia may coordinate these responses through a common molecular mechanism. Since hypoxia largely signals through stabilization of HIFs, we next examined the relative contribution of HIF-1α and HIF-2α to these effector programs. To test this, dLck^Cre^ and Hif1a^fl/fl^ or Hif2a^fl/fl^ mice were crossed to generate the deletion of HIF in T cells after positive selection in the thymus, and we refer to these mice as HIF-1a^cKO^ or HIF-2a^cKO^ mice. dLck^Cre^, Hif2a^fl/fl^, and Hif2a^fl/fl^ mice were used as controls (WT). Splenocytes or dLN cells from infected mice were stimulated in vitro with plate-bound anti-CD3 and soluble anti-CD28 under normoxic (21% O□) or hypoxic (1% O□) conditions for 48 hours. During the last 4 hours, cells also received PMA, ionomycin, and brefeldin A. Deficiency of HIF-1α significantly impaired hypoxia-induced granzyme B (Fig. 2A) and perforin expression (Fig. 2B). In contrast, HIF-2α deficiency had only a modest effect on the production of granzyme B and perforin (Fig. 2A-B).

**Figure 2.**
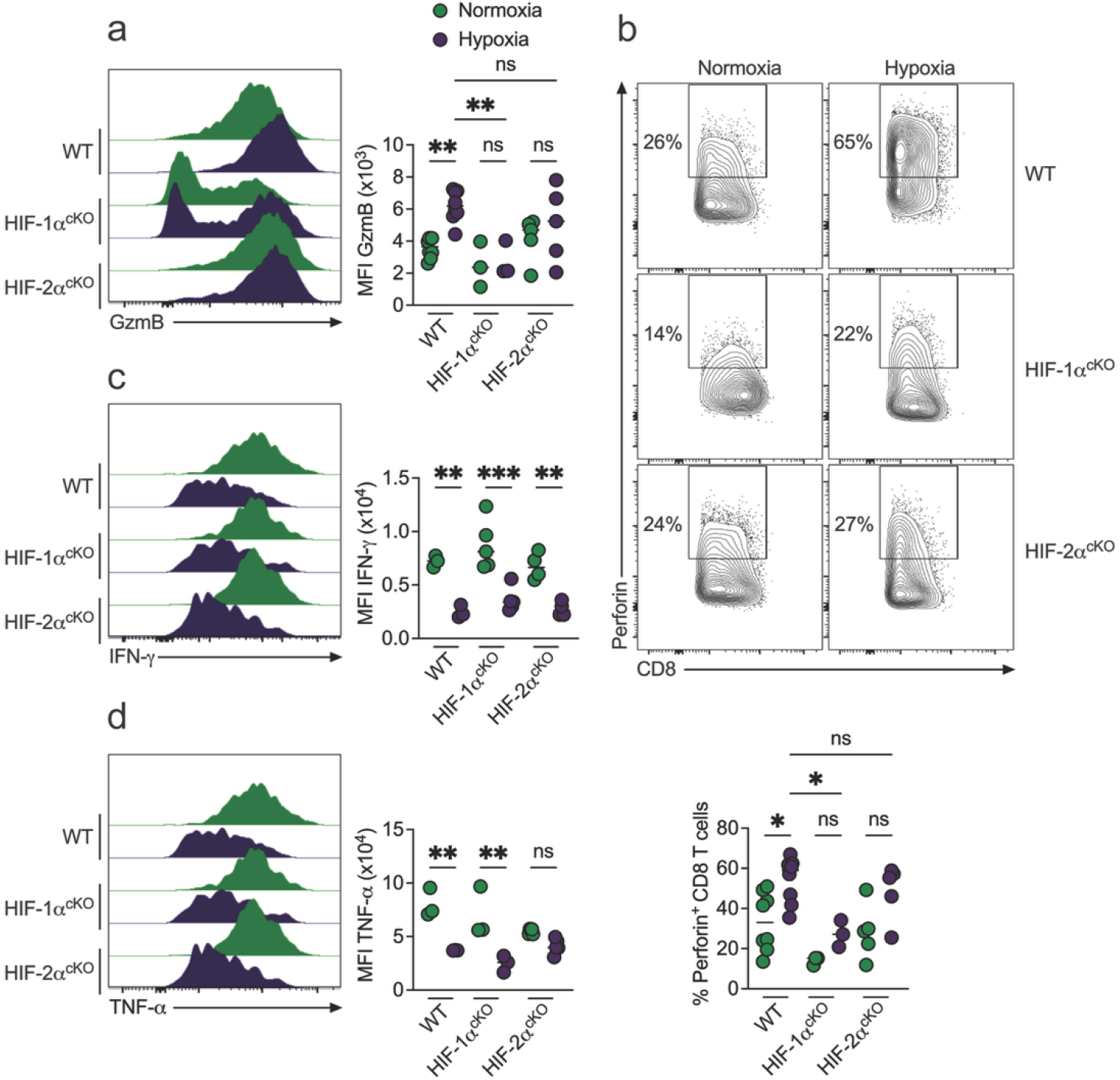
HIF-1α is required for hypoxic induction of granzyme B and perforin in CD8^+^ T cells. **(a and b)** dLN single cell suspensions of *Leishmania major*-infected wild-type (WT), HIF-1α^cKO^, and HIF-2α^cKO^ mice were incubated in normoxia (21% O_2_) or hypoxia (1% O_2_) for 48 hours with anti-CD3 and anti-CD28 stimulation and then CD8^+^ T cells were analyzed by flow cytometry for **(a)** mean fluorescence intensity (MFI) of granzyme B (GzmB) and **(b)** frequency of perforin expression. **(c and d)** Splenic single cell suspensions from *L. major*-infected wild type (WT), HIF-1α^cKO^, and HIF-2α^cKO^ mice were incubated in normoxia (21% O_2_) or hypoxia (1% O_2_) for 48 hours with anti-CD3 and anti-CD28 stimulation. For the last four hours of incubation, cells were stimulated with PMA (phorbol 12-myristate 13-acetate), ionomycin, and brefeldin A, and then CD8^+^ T cells were analyzed by flow cytometry for MFI of **(c)** IFN-γ and **(b)** TNF-α. Representative contour plots, histograms, and scatter dot plots showing individual mice are representative of 2 experiments with at least 3 mice per experiment. Gating strategy: live, single cells, CD45, CD3 or CD90, CD8β^+^, CD44^high^. ns=not significant, ∗P ≤ .05, ∗∗P ≤ .01, and ∗∗∗P ≤ .001 by 1-way ANOVA.

We next examined whether the suppression of protective cytokines was regulated through the same pathway. Deletion of either HIF-1α or HIF-2α failed to restore IFN-γ production under hypoxic conditions (Fig. 2C). Similarly, hypoxia suppressed TNF-α production in HIF-deficient CD8□ T cells (Fig. 2D). Together, these findings demonstrate that hypoxia regulates pathogenic and protective CD8□ T cell functions through distinct pathways. While HIF-1α signaling is required for induction of the cytotoxic program, suppression of type 1 cytokine production occurs independently of HIF.

### HIF-1α induces Blimp-1 expression

We previously identified the transcription factor Blimp-1 (encoded by the *Prdm1* gene) as a critical mediator of hypoxia-induced granzyme B expression, although perforin was induced through a Blimp-1-independent mechanism (8, 9). To explain the dual requirement for HIF-1α and Blimp-1 in granzyme B expression, we next investigated whether HIF signaling functions upstream of Blimp-1. Purified CD8□ T cells from infected WT, HIF-1α^cKO^, and HIF-2α^cKO^ mice were cultured under normoxic (21% O□) or hypoxic (1% O□) conditions, and expression of *Prdm1* was quantified after 24 hours by qPCR. Hypoxia robustly induced *Prdm1* expression in WT and HIF-2α-deficient CD8□ T cells. In contrast, HIF-1α-deficient CD8□ T cells failed to upregulate *Prdm1* following hypoxic stimulation (Fig. 3A). These data identify HIF-1α as an upstream regulator of Blimp-1 during acute hypoxic stimulation and elucidate the induction of granzyme B expression by HIF-1α.

**Figure 3.**
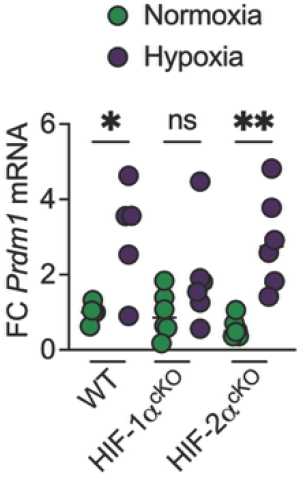
HIF-1α induces *Prdm1* expression in hypoxic CD8^+^ T cells. Isolated CD8^+^ T cells from the dLNs of *Leishmania major*-infected wild-type (WT), HIF-1α^cKO^, and HIF-2α^cKO^ mice were incubated in normoxia (21% O_2_) or hypoxia (1% O_2_) for 24 hours with anti-CD3, anti-CD28, and IL-2. Fold change (FC) of *Prdm1* mRNA over the average expression of normoxic cells was measured by qRT-PCR. Scatter dot plot showing individual mice combined from 2 independent experiments with at least 2 mice per experiment. ns=not significant, ∗P ≤ .05, and ∗∗P ≤ .01 by 1-way ANOVA.

### Hypoxia promotes an inhibitory phenotype without inducing terminal exhaustion

Because hypoxia suppressed production of protective cytokines independently of HIF signaling, we next asked whether hypoxia also changed the expression of inhibitory receptors associated with dysfunctional T cell responses. dLN cells from infected mice were stimulated in vitro with plate-bound anti-CD3 and soluble anti-CD28 under normoxic (21% O□) or hypoxic (1% O□) conditions for 48 hours. Exposure of activated CD8□ T cells to hypoxia significantly increased expression of PD-1, LAG-3, and TIM-3 (Fig. 4A-C), consistent with previous reports in chronic viral infection and cancer (18–20, 23, 26, 33, 36–41). Thus, hypoxia promotes not only cytotoxic differentiation but also acquisition of an inhibitory phenotype. Since sustained expression of inhibitory receptors is associated with terminal T cell exhaustion, we next examined expression of the lineage-defining transcription factor TOX in CD8□ T cells in lesions of mice infected with *Leishmania* (45, 46). Surprisingly, despite PD-1 expression, the vast majority of lesional CD8□ T cells remained TOX^low^, with only a small fraction exhibiting the PD-1^high^TOX^high^ phenotype characteristic of terminally exhausted cells (Fig. 4D). Together, these findings indicate that hypoxia promotes expression of inhibitory receptors but does not drive terminal exhaustion of lesional CD8□ T cells.

**Figure 4.**
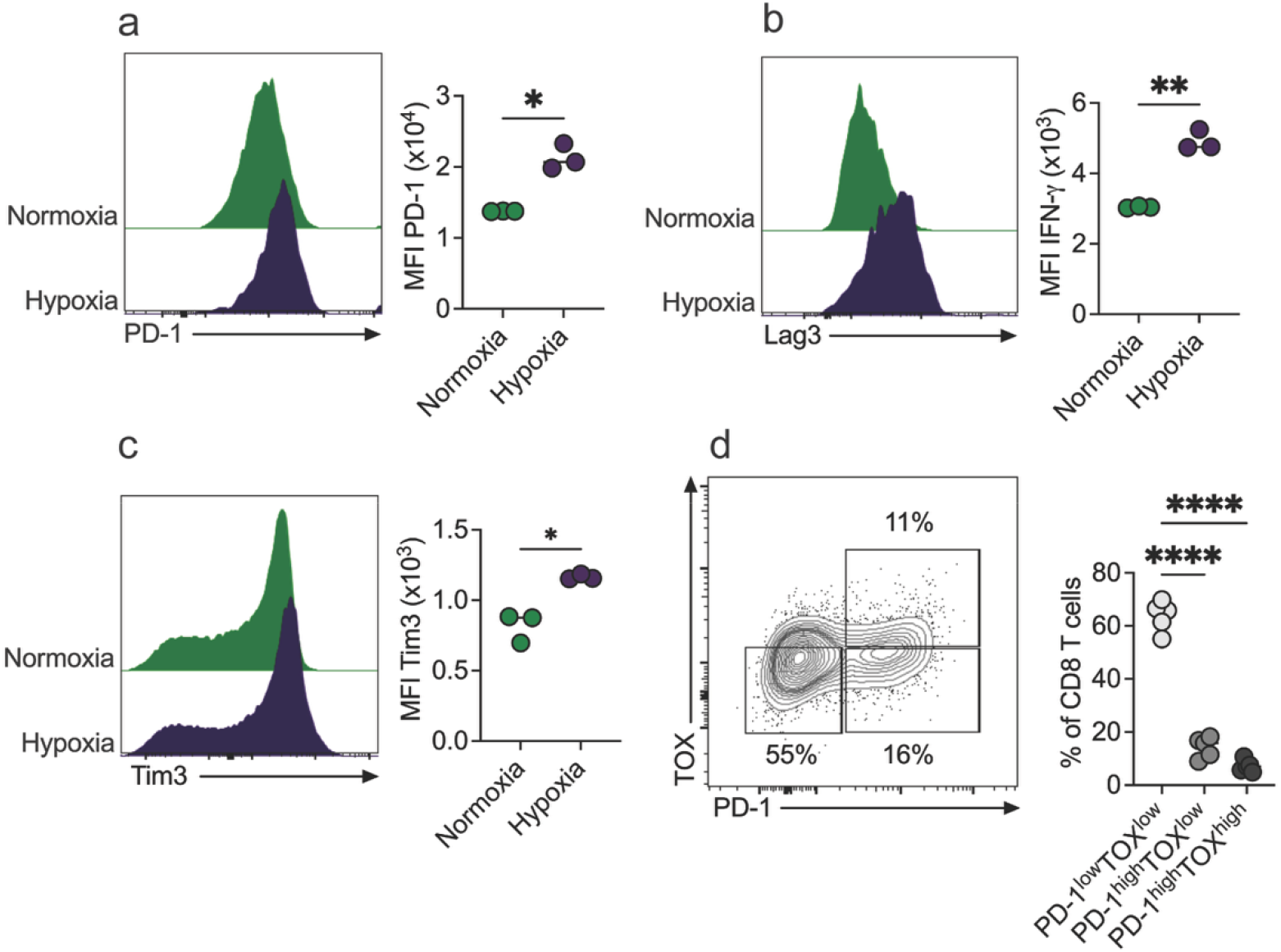
Hypoxia induces checkpoint molecules in CD8^+^ T cells. **(a-c)** dLN single-cell suspensions of *Leishmania major*-infected C57BL/6 mice were incubated in normoxia (21% O_2_) or hypoxia (1% O_2_) for 48 hours with anti-CD3 and anti-CD28 stimulation, and then CD8^+^ T cells were analyzed by flow cytometry for mean fluorescence intensity (MFI) of **(a)** PD-1, **(b)** Lag3, and **(c)** Tim3. **(d)** C57BL/6 mice were infected with *L. major*, and two to three weeks later, the lesions were analyzed by flow cytometry. The frequency of lesional CD8^+^ T cells was stratified into PD-1^low^TOX^low^, PD-1^high^TOX^low^, or PD-1^high^TOX^high^ groups. **(a-c)** Representative histograms and scatter dot plots showing individual mice are representative of 2 experiments with at least 3 mice per experiment. **(d)** Representative contour and scatter dot plot showing individual mice are representative of 3 experiments with at least 3 mice per experiment. (**a-c**) Gating strategy: live, single cells, CD90, CD8β^+^, CD44^high^. (**d**) Gating strategy: live, single cells, CD45, CD3, CD8β^+^, CD44^high^. ns=not significant, ∗P ≤ .05, ∗∗P ≤ .01, and ∗∗∗∗P ≤ .0001 by **(a-c)** 2-tailed Student’s t test or **(d)** by 1-way ANOVA.

### Both HIF-1α and HIF-2α contribute to CD8 T cell-mediated pathology during chronic infection

Our in vitro studies identified HIF-1α as an important regulator of acute hypoxia-induced cytotoxic differentiation. We next asked whether this relationship was maintained during chronic infection, where CD8□ T cells experience prolonged exposure to the hypoxic tissue microenvironment. To determine the cell-intrinsic contribution of HIF signaling to disease pathogenesis, Rag1^-/-^ mice were infected with *L. major* and immediately reconstituted with 3×10^6^ CD8□ T cells purified from the spleens of naïve WT, HIF-1α^cKO^, or HIF-2α^cKO^ mice (2, 3, 8, 9, 47, 48). As expected, transfer of WT CD8□ T cells resulted in severe progressive skin pathology (Fig. 5A-B). In contrast, mice receiving HIF-1α-deficient CD8□ T cells developed significantly reduced pathology, confirming an important role for HIF-1α in pathogenic CD8□ T cell function (Fig. 5A-B). Unexpectedly, HIF-2α deficiency also significantly reduced lesion size and pathology, despite having only a modest impact on acute hypoxic differentiation in vitro (Fig. 5A-B). Importantly, parasite burdens were unchanged among all experimental groups (Fig. 5C), consistent with our previous findings that lesional CD8□ T cells drive tissue pathology rather than parasite clearance. Taken together, these findings reveal an important distinction between acute and chronic hypoxic responses. While HIF-1α predominates during acute hypoxic induction of the cytotoxic program in vitro, both HIF-1α and HIF-2α contribute to CD8□ T cell-mediated pathology during chronic infection, suggesting complementary roles for these transcription factors within the chronically hypoxic tissue microenvironment.

**Figure 5.**
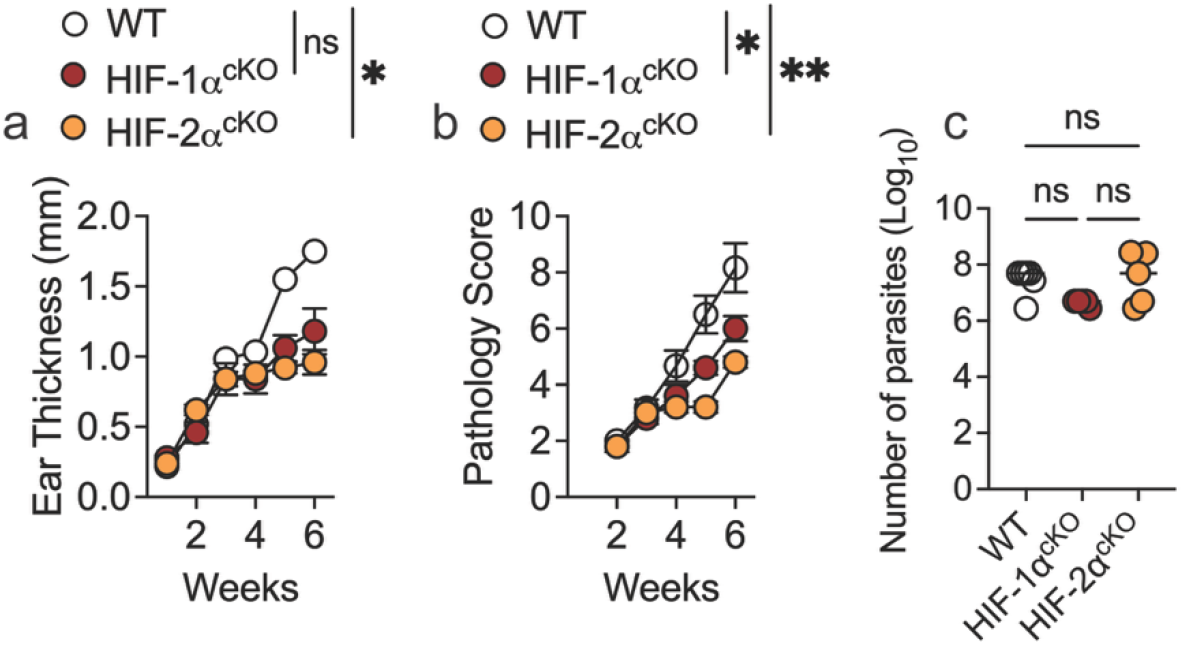
HIF-1α and HIF-2α contribute to CD8^+^ T cell-mediated pathology in cutaneous leishmaniasis. RAG^−/−^ mice were infected with *Leishmania major* and reconstituted with CD8^+^ T cells purified from wild-type (WT), HIF-1α ^cKO^, or HIF-2α ^cKO^ mice. **(a)** Ear thickness and **(b)** pathology score were measured weekly. **(b-c)** Statistical significance was determined at 5 weeks post-infection by a 2-tailed Student’s t-test compared to mice reconstituted with WT CD8^+^ T cells. ns=not significant, ∗P ≤ .05, and ∗∗P ≤ .01. **(d)** Parasite numbers at 6 weeks post-infection. ns=not significant by 1-way ANOVA. Data representative of 2 independent experiments with at least 3 mice per group.

## Discussion

The hypoxic microenvironment is increasingly recognized as an important regulator of immune responses in chronic infection, cancer, and inflammatory disease (5, 6). Although hypoxia is generally thought to enhance CD8□ T-cell effector function through activation of HIF signaling (18, 19, 24–26), our findings demonstrate that prolonged exposure to the chronically hypoxic environment of cutaneous leishmaniasis lesions has an unexpected effect. We show that hypoxia simultaneously enhances cytotoxicity and inhibitory receptor expression while suppressing the production of IFN-γ and TNF-α by CD8□ T cells. Surprisingly, these programs are regulated through distinct mechanisms. While HIF signaling was required for the induction of cytotoxic molecules and the pathogenic activity of lesional CD8O T cells, suppression of type 1 cytokine production occurred independently of HIF. These findings identify oxygen availability as a rheostat that uncouples protective and pathogenic CD8O T cell functions and reveal that distinct oxygen-sensing pathways independently control these effector programs.

The ability of hypoxia to separate cytotoxicity from cytokine production has important implications for our understanding of immunity during chronic *Leishmania* infection. In most settings, cytotoxicity and IFN-γ production are considered coordinated features of effector CD8□ T cells. Our data instead suggest that the chronically hypoxic tissue microenvironment preserves pathways responsible for tissue damage while limiting those that contribute to parasite control. The preferential preservation of cytotoxicity despite impaired cytokine production may represent an adaptive response to prolonged hypoxia. Because IFN-γ production is highly dependent on sustained glycolytic and biosynthetic activity, whereas cytotoxic function can rely, at least initially, on the release of preformed lytic granules, hypoxia may selectively restrict cytokine production while preserving the capacity to kill target cells (49–54). Alternatively, oxygen may directly regulate distinct transcriptional or epigenetic programs governing these effector functions (27, 34, 55–62). Regardless of the mechanism, these findings suggest that oxygen availability actively determines the balance between protective and pathogenic CD8O T cells within cutaneous leishmaniasis lesions.

Our findings also refine the relationship between hypoxia and T cell exhaustion. Previous studies in chronic viral infection and cancer have demonstrated that hypoxia and HIF signaling promote expression of inhibitory receptors without necessarily inducing terminal exhaustion (18, 19, 26). Consistent with these observations, hypoxia increased the expression of multiple inhibitory receptors in CD8O T cells, yet only a small subset of lesional cells acquired the canonical PD-1^high^TOX^high^ phenotype associated with terminal exhaustion. Together with our previous demonstration that hypoxia induces Blimp-1-dependent cytotoxic differentiation (8), these findings support the concept that chronic hypoxia promotes a unique functional adaptation characterized by preserved cytotoxicity despite acquisition of several checkpoint molecules. The signals that prevent progression toward terminal exhaustion remain unknown but likely reflect the unique inflammatory and metabolic environment of cutaneous leishmaniasis lesions.

The impact of hypoxia on T cell IFN-γ production remains controversial, with studies reporting both enhanced and diminished IFN-γ production, likely due to differences in acute versus chronic stimulation, HIF manipulation versus oxygen deprivation, and distinct disease settings (18–20, 24, 27–35). The HIF-independent suppression of IFN-γ observed here may be explained by previously described mechanisms limiting type 1 immunity during leishmaniasis. In visceral leishmaniasis, HIF-1α stabilization in dendritic cells suppresses IL-12 production and indirectly reduces IFN-γ expression by CD4□ T cells (63). In cutaneous leishmaniasis, IL-12 is present at relatively low levels within lesions, and previous studies demonstrated that exogenous IL-12 preferentially restores IFN-γ production by lesional CD8□ T cells rather than CD4□ Th1 cells (3). In addition to cell-extrinsic effects, HIF-independent cell-intrinsic effects might participate in suppressing IFN-γ. For example, oxygen-sensitive enzymes respond directly to oxygen availability independently of HIF. These include oxygen-dependent histone and DNA demethylases capable of altering chromatin accessibility and gene expression (27, 34, 55–62). Together, these observations suggest that chronic tissue hypoxia may suppress protective CD8□ T cell immunity through both direct effects on oxygen-sensitive signaling pathways and indirectly through regulation of antigen-presenting cell function.

The relative contributions of HIF-1α and HIF-2α appear to depend on the duration and context of hypoxic exposure. In vitro, HIF-1α deficiency more profoundly impaired granzyme B and perforin expression following acute hypoxic stimulation, whereas HIF-2α deficiency produced the strongest reduction in pathology in vivo. This observation is consistent with the distinct biology of the two HIF isoforms. HIF-1α is rapidly stabilized during acute reductions in oxygen and primarily regulates glycolytic adaptation, whereas HIF-2α accumulates during prolonged hypoxia and controls transcriptional programs involved in chronic tissue adaptation, angiogenesis, erythropoiesis, and iron metabolism (10–17). Because lesional CD8□ T cells reside within chronically hypoxic skin for several weeks, it is plausible that HIF-2α plays a more important role in sustaining pathogenic effector responses in vivo. Nevertheless, the overlapping functions of both isoforms suggest that maximal HIF activity within lesions likely results from the combined contributions of HIF-1α and HIF-2α.

Collectively, our study identifies chronic hypoxia as a dominant regulator of CD8O T cell differentiation within *Leishmania*-infected lesions and demonstrates that oxygen availability independently controls pathogenic and protective effector programs. Rather than inducing global T cell dysfunction, prolonged hypoxia selectively preserves cytotoxicity while suppressing type 1 cytokine production through distinct cell-intrinsic or -extrinsic mechanisms. These findings extend the concept that skin microenvironment actively determines immune cell function and suggest that therapeutic modulation of hypoxic signaling could rebalance protective and pathogenic CD8O T cell responses. Because chronic hypoxia is a defining feature of many infectious, inflammatory, and neoplastic diseases, these principles are likely to extend well beyond cutaneous leishmaniasis.

## Methods

### Mice

Our animal experiments were performed using male and female mice, and controls were sex matched, but sex was not specifically tested as a biological variable. C57BL/6 (6 weeks old) were purchased from Charles River, and dLCK^cre^ (Strain #:012837), HIF-1α ^fl/fl^ (Strain #:007561), HIF-2α ^fl/fl^ (Strain #:008407), and Rag^-/-^ (Strain #:012212216) were purchased from The Jackson Laboratory and bred in our mouse facility. dLCK^cre^ and HIF-1α ^fl/fl^, or HIF-2α ^fl/fl^, mice were crossed to generate the deletion of HIF-1α or HIF-2α in T cells after positive selection in the thymus and were referred to as HIF-1α ^cKO^ or HIF-2α ^cKO^ respectively. dLCK^cre^, HIF-1α ^fl/fl^, or HIF-2α ^fl/fl^ mice were used as wild type (WT) controls. All mice were maintained in a specific pathogen-free environment at The Ohio State University

### Parasites

*Leishmania major* (strain WHO/MHOM/IL/80/Friedlin) were grown in Schneider’s insect medium (GIBCO) supplemented with 20% heat-inactivated FBS (Sigma) and 2 mM glutamine (Thermo-fisher). Metacyclic-enriched promastigotes were used for infection (64). Mice were infected with 10^6^ *L. major* intradermally in the ear, and the lesion progression was monitored weekly by measuring the ear thickness with a digital caliper (Fisher Scientific). A standardized pathology score (ranging from 0 to 12) was used to quantify macroscopic lesion progression, disease severity, tissue damage, and inflammation in infected mice. Scores for redness, deformation of the ear pinnae, ulceration, and tissue loss varied from no symptom (0), mild (1), moderate (2), and severe (3) for each individual mouse.

### Cell purification and adoptive transfer

For the Rag1^−/−^ mouse model, splenocytes from WT, HIF-1α ^cKO^, and HIF-2α ^cKO^ mice were collected, RBCs were lysed with ACK lysing buffer (Lonza), and CD8^+^ T cells were purified using a magnetic bead separation kit (Miltenyi Biotec). CD8^+^ T cells (3×10^6^ cells) were transferred intravenously into Rag1^−/−^ mice that were subsequently infected with *L. major*. Mice reconstituted with CD8^+^ T cells received 4 injections of 250 μg anti-CD4 (clone GK1.5, Bio X Cell, catalog BE0003-1) within the first 2 weeks

### Single-cell suspension preparation

Infected ears were harvested, the dorsal and ventral layers of the ear separated, and the ears were incubated in RPMI 1640 (Gibco) with 250 μg/mL of Liberase TL (Roche Diagnostics) and DNAse I (Sigma) and dissociated using the gentleMACS™ Dissociator with Heaters (Miltenyi Biotec) using program 37_Multi_H. An aliquot of the cell suspension was used for parasite titration for each tissue. Draining lymph nodes (dLNs) were homogenized using a cell strainer (40 μm, BD Pharmingen) to obtain single-cell suspensions. Spleens were homogenized using a cell strainer (40 μm, BD Pharmingen) to obtain single-cell suspensions, and red blood cells were lysed with ACK lysis buffer (Lonza). For analysis of IFN-γ or TNF-α, single-cell suspensions were stimulated for four hours with 0.2 ug/mL phorbol 12-myristate 13-acetate (PMA; Sigma), 2 ug/mL ionomycin (Sigma), and 5 ug/mL of Brefeldin A (BioLegend).

### Parasite titration

The parasite burden in the ears was quantified as described previously (65). Briefly, the homogenate was serially diluted and incubated at 26°C. The number of viable parasites was calculated from the highest dilution at which parasites were observed after 7 days

### In vitro stimulation

For flow cytometry analysis, dLN or spleen single-cell suspensions from infected mice plated into 48 well plates “U” bottom at a concentration of 0.5×10^6^ cells/ml with 5 μg/mL of plate-bound anti-CD3 (clone 145-2C11, Invitrogen), and 0.5 μg/mL soluble anti-CD28 (clone 37.51, Invitrogen) at 37°C and 5% CO_2_ for 48 hours in RPMI 1640 containing 100 Units of penicillin and 0.1 mg/mL of Streptomycin (Sigma), 2mM L-Glutamine (Thermo-fisher) and 10% FBS (Sigma). Cells were cultured under normoxic conditions (standard incubator, 21% oxygen) or 1% oxygen in either a Modulating Incubator Chamber (Billups-Rothenberg) or Baker Ruskin InvivO_2_ 400, an incubator equipped to replace oxygen with nitrogen. For analysis of IFN-γ or TNF-α, cells were stimulated for four hours with 0.2 ug/mL phorbol 12-myristate 13-acetate (PMA; Sigma), 2 ug/mL ionomycin (Sigma), and 5 ug/mL of Brefeldin A (BioLegend).

For mRNA isolation, dLN single-cell suspensions from infected mice were obtained and CD8^+^ T cells were purified using a magnetic bead separation kit (Miltenyi Biotec) then plated into 48 well plates “U” bottom at a concentration of 1×10^6^ cells/ml with 5 μg/mL of plate-bound anti-CD3 (clone 145-2C11, Invitrogen), and 0.5 μg/mL soluble anti-CD28 (clone 37.51, Invitrogen), and 20 unit/mL of IL-2 (Hoffman La-Roche, Ro 23-6019) at 37°C and 5% CO_2_ for 24 hours in RPMI 1640 containing 100 Units of penicillin and 0.1 mg/mL of Streptomycin (Sigma), 2mM L-Glutamine (Thermo-fisher) and 10% FBS (Sigma). Cells were cultured under normoxic or hypoxic conditions as described above.

### Quantification of mRNA by qPCR

RNA extraction was performed using nulceospin RNA Mini kit (Macherery-nagel) and used to prepare cDNA using iScript Reverse Transcription Supermix for RT-qPCR (Bio-Rad Laboratories) on ABS Proflex Thermal cycler (Thermo Fisher Scientific). qPCR was carried out on a C1000 RT-PCR using iTaq Universal SYBR Green Supermix (all from Bio-Rad Laboratories) and primers targeting *Prdm1* (forward, 5′-TTCTCTTGGAAAAACGTGTG G-3′ and reverse, 5′-GGAGCCGGAGCTAGACTTG-3′). The qPCR results were normalized to *Actb* (forward 5′-CGC TGT ATT CCC CTC CAT CG-3′ and reverse 5′CCA GTT GGT AAC AAT GCC ATG T-3′). All reactions were carried out in duplicates, and data are represented as fold change over the average expression of normoxic cells.

### Flow cytometric analysis

Before surface and intracellular staining, cell suspensions were stained with either LIVE/DEAD fixable aqua dead cell stain kit (L34957), LIVE/DEAD fixable blue dead cell stain kit (L23105), or eBioscience Fixable Viability Dye eFluor-780 (65-0865-14, all Thermo-fisher) according to manufacturer instructions. Cell analysis was performed using the FlowJo Software (Tree Star), and gates were created based on fluorescence minus one control. The following antibodies were used: CD45 (clone 30-F11, catalog 63-0451-80), GzmB (clone GB11, catalog GRB05 or GRB04), CD3 (clone 17A2, catalog 48-0032-82), CD44 (clone IM7, catalog 12-0440-83), TOX (clone TXRX10, catalog 50-6502-82), IFN-γ (clone XMG1.2, catalog 25-7311-82), TIM-3 (clone RMT3-23, catalog 416-5870-82), and TNF-α (clone MP6-XT22, catalog 17-7321-82) (all from Invitrogen, Thermo Fisher Scientific); CD3 (clone 17A2, catalog 741319), and CD44 (clone IM7, catalog 751414) (all from BD Biosciences); CD45 (clone 30-F11, catalog 103137), CD8β (clone YTS156.7.7, catalog 126610 or 126633), LAG-3 (clone C9B7W, catalog 125206 or 125226), CD90 (clone 53-2.1, catalog 140306), PD-1 (clone 29F.1A12, catalog 135224, and perforin (clone S16009A, catalog 154304) (all from BioLegend). The stained cells were acquired on BD FACSymphony A3 (BD Biosciences), Cytek Aurora 4 Laser (UV-V-B-R), or Cytek Aurora 5 Laser (UV-V-B-YG-R).

### Statistics

For differences between two groups, statistical significance was determined using unpaired Student’s t-test. For multiple comparisons, one-way analysis of variance (ANOVA) was performed. Differences were considered significant when p ≤ 0.05 (*), p ≤ 0.01 (**), p ≤ 0.001 (***), or p ≤ 0.0001 (****) and ns = not significant.

### Study Approval

This study was carried out per the recommendations in the Guide for the Care and Use of Laboratory Animals of the National Institutes of Health. The Institutional Animal Care and Use Committee and The Ohio State University approved the protocol.

## Author contributions

E.A.F. and F.O.N. designed research studies; E.A.F., O.L.S., and F.S. conducted experiments; E.A.F. and F.O.N. analyzed and interpreted the data; E.A.F. wrote the manuscript; F.O.N. edited the manuscript and supervised the study.

## Bibliography

1. Scott, P., and F. O. Novais. 2016. Cutaneous leishmaniasis: immune responses in protection and pathogenesis. Nat Rev Immunol 16: 581–592.

2. Novais, F. O., L. P. Carvalho, J. W. Graff, D. P. Beiting, G. Ruthel, D. S. Roos, M. R. Betts, M. H. Goldschmidt, M. E. Wilson, C. I. de Oliveira, and P. Scott. 2013. Cytotoxic T Cells Mediate Pathology and Metastasis in Cutaneous Leishmaniasis. PLOS Pathogens 9: e1003504.

3. Novais, F. O., A. C. Wong, D. O. Villareal, D. P. Beiting, and P. Scott. 2018. CD8+ T Cells Lack Local Signals To Produce IFN-γ in the Skin during Leishmania Infection. J Immunol 200: 1737–1745.

4. Fowler, E. A., and F. O. Novais. 2025. Step-by-step: the CD8 T cell journey in leishmaniasis. mBio 16: e03537–23.

5. Mirchandani, A. S., M. A. Sanchez-Garcia, and S. R. Walmsley. 2025. How oxygenation shapes immune responses: emerging roles for physioxia and pathological hypoxia. Nat Rev Immunol 25: 161–177.

6. Lee, P., N. S. Chandel, and M. C. Simon. 2020. Cellular adaptation to hypoxia through hypoxia inducible factors and beyond. Nat Rev Mol Cell Biol 21: 268–283.

7. Mahnke, A., R. J. Meier, V. Schatz, J. Hofmann, K. Castiglione, U. Schleicher, O. S. Wolfbeis, C. Bogdan, and J. Jantsch. 2014. Hypoxia in *Leishmania major* Skin Lesions Impairs the NO-Dependent Leishmanicidal Activity of Macrophages. Journal of Investigative Dermatology 134: 2339–2346.

8. Fowler, E. A., C. F. Amorim, K. Mostacada, A. Yan, L. A. Sacramento, R. A. Stanco, E. D. S. Hales, A. Varkey, W. Zong, G. D. Wu, C. I. de Oliveira, P. L. Collins, and F. O. Novais. 2024. Neutrophil-mediated hypoxia drives pathogenic CD8^+^ T cell responses in cutaneous leishmaniasis. J Clin Invest 134.

9. Fowler, E. A., L. Amorim Sacramento, B. A. Bowman, B. Lee, C.-W. J. Lio, Y.-D. Dong, J. A. Spicer, J. A. Trapani, and F. O. Novais. 2025. Hypoxia and IL-15 Cooperate to Induce Perforin Expression by CD8 T Cells and Promote Damage to the Skin in Murine Cutaneous Leishmaniasis. Journal of Investigative Dermatology 145: 3145–3157.e1.

10. Semenza, G. L. 1994. Regulation of Erythropoietin Production: New Insights Into Molecular Mechanisms of Oxygen Homeostasis. Hematology/Oncology Clinics of North America 8: 863–884.

11. Ebert, B. L., J. D. Firth, and P. J. Ratcliffe. 1995. Hypoxia and Mitochondrial Inhibitors Regulate Expression of Glucose Transporter-1 via Distinct Cis-acting Sequences (*). Journal of Biological Chemistry 270: 29083–29089.

12. Semenza, G. L., B.-H. Jiang, S. W. Leung, R. Passantino, J.-P. Concordet, P. Maire, and A. Giallongo. 1996. Hypoxia Response Elements in the Aldolase A, Enolase 1, and Lactate Dehydrogenase A Gene Promoters Contain Essential Binding Sites for Hypoxia-inducible Factor 1*. Journal of Biological Chemistry 271: 32529–32537.

13. Hu, C.-J., L.-Y. Wang, L. A. Chodosh, B. Keith, and M. C. Simon. 2003. Differential Roles of Hypoxia-Inducible Factor 1α (HIF-1α) and HIF-2α in Hypoxic Gene Regulation. Molecular and Cellular Biology 23: 9361–9374.

14. Papandreou, I., R. A. Cairns, L. Fontana, A. L. Lim, and N. C. Denko. 2006. HIF-1 mediates adaptation to hypoxia by actively downregulating mitochondrial oxygen consumption. Cell Metabolism 3: 187–197.

15. Rankin, E. B., M. P. Biju, Q. Liu, T. L. Unger, J. Rha, R. S. Johnson, M. C. Simon, B. Keith, and V. H. Haase. 2007. Hypoxia-inducible factor–2 (HIF-2) regulates hepatic erythropoietin in vivo. J Clin Invest 117: 1068–1077.

16. Koh, M. Y., R. Lemos Jr, X. Liu, and G. Powis. 2011. The Hypoxia-Associated Factor Switches Cells from HIF-1α- to HIF-2α-Dependent Signaling Promoting Stem Cell Characteristics, Aggressive Tumor Growth and Invasion. Cancer Res 71: 4015–4027.

17. Bowman, B. A., and F. O. Novais. 2025. Oxygen and immunity to Leishmania infection. Infection and Immunity 93: e00504–24.

18. Doedens, A. L., A. T. Phan, M. H. Stradner, J. K. Fujimoto, J. V. Nguyen, E. Yang, R. S. Johnson, and A. W. Goldrath. 2013. Hypoxia-inducible factors enhance the effector responses of CD8+ T cells to persistent antigen. Nat Immunol 14: 1173–1182.

19. Palazon, A., P. A. Tyrakis, D. Macias, P. Veliça, H. Rundqvist, S. Fitzpatrick, N. Vojnovic, A. T. Phan, N. Loman, I. Hedenfalk, T. Hatschek, J. Lövrot, T. Foukakis, A. W. Goldrath, J. Bergh, and R. S. Johnson. 2017. An HIF-1α/VEGF-A Axis in Cytotoxic T Cells Regulates Tumor Progression. Cancer Cell 32: 669–683.e5.

20. Scharping, N. E., D. B. Rivadeneira, A. V. Menk, P. D. A. Vignali, B. R. Ford, N. L. Rittenhouse, R. Peralta, Y. Wang, Y. Wang, K. DePeaux, A. C. Poholek, and G. M. Delgoffe. 2021. Mitochondrial stress induced by continuous stimulation under hypoxia rapidly drives T cell exhaustion. Nat Immunol 22: 205–215.

21. McNamee, E. N., D. Korns Johnson, D. Homann, and E. T. Clambey. 2013. Hypoxia and hypoxia-inducible factors as regulators of T cell development, differentiation, and function. Immunol Res 55: 58–70.

22. Phan, A. T., and A. W. Goldrath. 2015. Hypoxia-inducible factors regulate T cell metabolism and function. Molecular Immunology 68: 527–535.

23. Wu, H., X. Zhao, S. M. Hochrein, M. Eckstein, G. F. Gubert, K. Knöpper, A. M. Mansilla, A. Öner, R. Doucet-Ladevèze, W. Schmitz, B. Ghesquière, S. Theurich, J. Dudek, G. Gasteiger, A. Zernecke, S. Kobold, W. Kastenmüller, and M. Vaeth. 2023. Mitochondrial dysfunction promotes the transition of precursor to terminally exhausted T cells through HIF-1α-mediated glycolytic reprogramming. Nat Commun 14: 6858.

24. Finlay, D. K., E. Rosenzweig, L. V. Sinclair, C. Feijoo-Carnero, J. L. Hukelmann, J. Rolf, A. A. Panteleyev, K. Okkenhaug, and D. A. Cantrell. 2012. PDK1 regulation of mTOR and hypoxia-inducible factor 1 integrate metabolism and migration of CD8+ T cells. J Exp Med 209: 2441–2453.

25. Veliça, P., P. P. Cunha, N. Vojnovic, I. P. Foskolou, D. Bargiela, M. Gojkovic, H. Rundqvist, and R. S. Johnson. 2021. Modified Hypoxia-Inducible Factor Expression in CD8+ T Cells Increases Antitumor Efficacy. Cancer Immunol Res 9: 401–414.

26. Liikanen, I., C. Lauhan, S. Quon, K. Omilusik, A. T. Phan, L. B. Bartrolí, A. Ferry, J. Goulding, J. Chen, J. P. Scott-Browne, J. T. Yustein, N. E. Scharping, D. A. Witherden, and A. W. Goldrath. 2021. Hypoxia-inducible factor activity promotes antitumor effector function and tissue residency by CD8^+^ T cells. J Clin Invest 131.

27. Cunha, P. P., E. Minogue, L. C. Krause, R. M. Hess, D. Bargiela, B. J. Wadsworth, L. Barbieri, C. Brombach, I. P. Foskolou, I. Bogeski, P. Velica, and R. S. Johnson. 2023. Oxygen levels at the time of activation determine T cell persistence and immunotherapeutic efficacy. eLife 12: e84280.

28. de Almeida, P. E., J. Mak, G. Hernandez, R. Jesudason, A. Herault, V. Javinal, J. Borneo, J. M. Kim, and K. B. Walsh. 2020. Anti-VEGF Treatment Enhances CD8+ T-cell Antitumor Activity by Amplifying Hypoxia. Cancer Immunol Res 8: 806–818.

29. Vuillefroy de Silly, R., L. Ducimetière, C. Yacoub Maroun, P.-Y. Dietrich, M. Derouazi, and P. R. Walker. 2015. Phenotypic switch of CD8+ T cells reactivated under hypoxia toward IL-10 secreting, poorly proliferative effector cells. European Journal of Immunology 45: 2263–2275.

30. Chen, P.-M., P. C. Wilson, J. A. Shyer, M. Veselits, H. R. Steach, C. Cui, G. Moeckel, M. R. Clark, and J. Craft. 2020. Kidney tissue hypoxia dictates T cell–mediated injury in murine lupus nephritis. Science Translational Medicine 12: eaay1620.

31. Caldwell, C. C., H. Kojima, D. Lukashev, J. Armstrong, M. Farber, S. G. Apasov, and M. V. Sitkovsky. 2001. Differential Effects of Physiologically Relevant Hypoxic Conditions on T Lymphocyte Development and Effector Functions. J Immunol 167: 6140–6149.

32. Murthy, A., S. A. Gerber, C. J. Koch, and E. M. Lord. 2019. Intratumoral Hypoxia Reduces IFN-γ–Mediated Immunity and MHC Class I Induction in a Preclinical Tumor Model. Immunohorizons 3: 149–160.

33. Ross, S. H., C. M. Rollings, and D. A. Cantrell. 2021. Quantitative Analyses Reveal How Hypoxia Reconfigures the Proteome of Primary Cytotoxic T Lymphocytes. Front. Immunol. 12.

34. Ma, S., Y. Zhao, W. C. Lee, L.-T. Ong, P. L. Lee, Z. Jiang, G. Oguz, Z. Niu, M. Liu, J. Y. Goh, W. Wang, M. A. Bustos, S. Ehmsen, A. Ramasamy, D. S. B. Hoon, H. J. Ditzel, E. Y. Tan, Q. Chen, and Q. Yu. 2022. Hypoxia induces HIF1α-dependent epigenetic vulnerability in triple negative breast cancer to confer immune effector dysfunction and resistance to anti-PD-1 immunotherapy. Nat Commun 13: 4118.

35. Shen, H., O. A. Ojo, H. Ding, L. J. Mullen, C. Xing, M. I. Hossain, A. Yassin, V. Y. Shi, Z. Lewis, E. Podgorska, S. A. Andrabi, M. R. Antoniewicz, J. A. Bonner, and L. Z. Shi. 2024. HIF1α-regulated glycolysis promotes activation-induced cell death and IFN-γ induction in hypoxic T cells. Nat Commun 15: 9394.

36. Vignali, P. D. A., K. DePeaux, M. J. Watson, C. Ye, B. R. Ford, K. Lontos, N. K. McGaa, N. E. Scharping, A. V. Menk, S. C. Robson, A. C. Poholek, D. B. Rivadeneira, and G. M. Delgoffe. 2023. Hypoxia drives CD39-dependent suppressor function in exhausted T cells to limit antitumor immunity. Nat Immunol 24: 267–279.

37. Baessler, A., and D. A. A. Vignali. 2024. T Cell Exhaustion. Annual Review of Immunology 42: 179–206.

38. Kim, A.-R., S. J. Choi, J. Park, M. Kwon, T. Chowdhury, H. J. Yu, S. Kim, H. Kang, K.-M. Kim, S.-H. Park, C.-K. Park, and E.-C. Shin. 2022. Spatial immune heterogeneity of hypoxia-induced exhausted features in high-grade glioma. OncoImmunology 11: 2026019.

39. Liu, Y.-N., J.-F. Yang, D.-J. Huang, H.-H. Ni, C.-X. Zhang, L. Zhang, J. He, J.-M. Gu, H.-X. Chen, H.-Q. Mai, Q.-Y. Chen, X.-S. Zhang, S. Gao, and J. Li. 2020. Hypoxia Induces Mitochondrial Defect That Promotes T Cell Exhaustion in Tumor Microenvironment Through MYC-Regulated Pathways. Front. Immunol. 11.

40. Sun, Q., and C. Dong. 2026. Regulators of CD8+ T cell exhaustion. Nat Rev Immunol 26: 129–151.

41. Franco, F., A. Jaccard, P. Romero, Y.-R. Yu, and P.-C. Ho. 2020. Metabolic and epigenetic regulation of T-cell exhaustion. Nat Metab 2: 1001–1012.

42. da Fonseca-Martins, A. M., T. D. Ramos, J. E. S. Pratti, L. Firmino-Cruz, D. C. O. Gomes, L. Soong, E. M. Saraiva, and H. L. de Matos Guedes. 2019. Immunotherapy using anti-PD-1 and anti-PD-L1 in Leishmania amazonensis-infected BALB/c mice reduce parasite load. Sci Rep 9: 20275.

43. Freitas e Silva, R. de, R. I. Gálvez, V. R. A. Pereira, M. E. F. de Brito, S. L. Choy, H. Lotter, L. Bosurgi, and T. Jacobs. 2020. Programmed Cell Death Ligand (PD-L)-1 Contributes to the Regulation of CD4+ T Effector and Regulatory T Cells in Cutaneous Leishmaniasis. Front. Immunol. 11.

44. Garcia de Moura, R., L. P. Covre, C. H. Fantecelle, V. A. T. Gajardo, C. B. Cunha, L. L. Stringari, A. T. Belew, C. B. Daniel, S. V. V. Zeidler, C. E. Tadokoro, H. L. de Matos Guedes, R. L. Zanotti, D. Mosser, A. Falqueto, A. N. Akbar, and D. C. O. Gomes. 2021. PD-1 Blockade Modulates Functional Activities of Exhausted-Like T Cell in Patients With Cutaneous Leishmaniasis. Front. Immunol. 12.

45. Khan, O., J. R. Giles, S. McDonald, S. Manne, S. F. Ngiow, K. P. Patel, M. T. Werner, A. C. Huang, K. A. Alexander, J. E. Wu, J. Attanasio, P. Yan, S. M. George, B. Bengsch, R. P. Staupe, G. Donahue, W. Xu, R. K. Amaravadi, X. Xu, G. C. Karakousis, T. C. Mitchell, L. M. Schuchter, J. Kaye, S. L. Berger, and E. J. Wherry. 2019. TOX transcriptionally and epigenetically programs CD8+ T cell exhaustion. Nature 571: 211–218.

46. Huang, Y. J., S. F. Ngiow, A. E. Baxter, S. Manne, S. L. Park, J. E. Wu, O. Khan, J. R. Giles, and E. J. Wherry. 2025. Continuous expression of TOX safeguards exhausted CD8 T cell epigenetic fate. Science Immunology 10: eado3032.

47. Novais, F. O., A. M. Carvalho, M. L. Clark, L. P. Carvalho, D. P. Beiting, I. E. Brodsky, E. M. Carvalho, and P. Scott. 2017. CD8+ T cell cytotoxicity mediates pathology in the skin by inflammasome activation and IL-1β production. PLOS Pathogens 13: e1006196.

48. Belkaid, Y., C. A. Piccirillo, S. Mendez, E. M. Shevach, and D. L. Sacks. 2002. CD4+CD25+ regulatory T cells control Leishmania major persistence and immunity. Nature 420: 502–507.

49. de Saint Basile, G., G. Ménasché, and A. Fischer. 2010. Molecular mechanisms of biogenesis and exocytosis of cytotoxic granules. Nat Rev Immunol 10: 568–579.

50. Voskoboinik, I., J. C. Whisstock, and J. A. Trapani. 2015. Perforin and granzymes: function, dysfunction and human pathology. Nat Rev Immunol 15: 388–400.

51. Ritter, A. T., Y. Asano, J. C. Stinchcombe, N. M. G. Dieckmann, B.-C. Chen, C. Gawden-Bone, S. van Engelenburg, W. Legant, L. Gao, M. W. Davidson, E. Betzig, J. Lippincott-Schwartz, and G. M. Griffiths. 2015. Actin Depletion Initiates Events Leading to Granule Secretion at the Immunological Synapse. Immunity 42: 864–876.

52. Cham, C. M., and T. F. Gajewski. 2005. Glucose Availability Regulates IFN-γ Production and p70S6 Kinase Activation in CD8+ Effector T Cells. J Immunol 174: 4670–4677.

53. Chang, C.-H., J. D. Curtis, L. B. Maggi, B. Faubert, A. V. Villarino, D. O’Sullivan, S. C.-C. Huang, G. J. W. van der Windt, J. Blagih, J. Qiu, J. D. Weber, E. J. Pearce, R. G. Jones, and E. L. Pearce. 2013. Posttranscriptional Control of T Cell Effector Function by Aerobic Glycolysis. Cell 153: 1239–1251.

54. Cham, C. M., G. Driessens, J. P. O’Keefe, and T. F. Gajewski. 2008. Glucose Deprivation Inhibits Multiple Key Gene Expression Events and Effector Functions in CD8+ T Cells. Eur J Immunol 38: 2438–2450.

55. Wang, F., R. Zhang, T. V. Beischlag, C. Muchardt, M. Yaniv, and O. Hankinson. 2004. Roles of Brahma and Brahma/SWI2-Related Gene 1 in Hypoxic Induction of the Erythropoietin Gene *. Journal of Biological Chemistry 279: 46733–46741.

56. Johnson, A. B., N. Denko, and M. C. Barton. 2008. Hypoxia induces a novel signature of chromatin modifications and global repression of transcription. Mutation Research - Fundamental and Molecular Mechanisms of Mutagenesis 640: 174–179.

57. Kenneth, N. S., S. Mudie, P. van Uden, and S. Rocha. 2009. SWI/SNF Regulates the Cellular Response to Hypoxia *. Journal of Biological Chemistry 284: 4123–4131.

58. Thienpont, B., J. Steinbacher, H. Zhao, F. D’Anna, A. Kuchnio, A. Ploumakis, B. Ghesquière, L. Van Dyck, B. Boeckx, L. Schoonjans, E. Hermans, F. Amant, V. N. Kristensen, K. P. Koh, M. Mazzone, M. L. Coleman, T. Carell, P. Carmeliet, and D. Lambrechts. 2016. Tumour hypoxia causes DNA hypermethylation by reducing TET activity. Nature 537: 63–68.

59. Xia, X., M. E. Lemieux, W. Li, J. S. Carroll, M. Brown, X. S. Liu, and A. L. Kung. 2009. Integrative analysis of HIF binding and transactivation reveals its role in maintaining histone methylation homeostasis. Proceedings of the National Academy of Sciences 106: 4260–4265.

60. Intlekofer, A. M., R. G. Dematteo, S. Venneti, L. W. S. Finley, C. Lu, A. R. Judkins, A. S. Rustenburg, P. B. Grinaway, J. D. Chodera, J. R. Cross, and C. B. Thompson. 2015. Hypoxia Induces Production of L-2-Hydroxyglutarate. Cell Metabolism 22: 304–311.

61. Batie, M., J. Frost, M. Frost, J. W. Wilson, P. Schofield, and S. Rocha. 2019. Hypoxia induces rapid changes to histone methylation and reprograms chromatin. Science 363: 1222–1226.

62. Chakraborty, A. A., T. Laukka, M. Myllykoski, A. E. Ringel, M. A. Booker, M. Y. Tolstorukov, Y. J. Meng, S. R. Meier, R. B. Jennings, A. L. Creech, Z. T. Herbert, S. K. McBrayer, B. A. Olenchock, J. D. Jaffe, M. C. Haigis, R. Beroukhim, S. Signoretti, P. Koivunen, and W. G. Kaelin. 2019. Histone demethylase KDM6A directly senses oxygen to control chromatin and cell fate. Science 363: 1217–1222.

63. Hammami, A., B. M. Abidin, K. M. Heinonen, and S. Stäger. 2018. HIF-1α hampers dendritic cell function and Th1 generation during chronic visceral leishmaniasis. Sci Rep 8: 3500.

64. Späth, G. F., and S. M. Beverley. 2001. A Lipophosphoglycan-Independent Method for Isolation of Infective *Leishmania* Metacyclic Promastigotes by Density Gradient Centrifugation. Experimental Parasitology 99: 97–103.

65. Uzonna, J. E., K. L. Joyce, and P. Scott. 2004. Low Dose Leishmania major Promotes a Transient T Helper Cell Type 2 Response That Is Down-regulated by Interferon γ–producing CD8+ T Cells. J Exp Med 199: 1559–1566.

